# Hyperthermic Seizure Susceptibility and Focal Decreases in Parvalbumin-Expressing Cortical Interneurons in a Mouse Model of PCDH19-Clustering Epilepsy

**DOI:** 10.1101/2025.11.12.688097

**Authors:** Julie M. Ziobro, Joy Huang, Noor Daddo, Emma Margolis, Kira Jonatzke, Hisashi Umemori, Jack M. Parent

## Abstract

**Objective:** Protocadherin-19 (PCDH19)-clustering epilepsy (PCE) is a severe genetic epilepsy that manifests with early-onset cluster seizures often triggered by fever, intellectual disability, autistic features, and later neuropsychiatric risk. *PCDH19* is an X-linked gene critical for brain development. PCE predominantly affects females and rare mosaic males, but not hemizygous males, likely due to cellular mosaicism arising from random X-inactivation and resultant segregation of wild-type and mutant neurons (so-called cellular interference) during development. We generated a novel PCE mouse model and explored neuronal segregation, seizure susceptibility and cortical interneuron distributions.

**Methods:** Female *Pcdh19^+/-^* mice were crossed with X-GFP males to visualize random X-inactivation patterns. Seizure susceptibility was assessed in juvenile mice using hyperthermia and flurothyl exposure. Behavioral testing evaluated cognitive domains. Interneuron distribution in hippocampus and cortex was examined histologically by immunolabeling and crosses with parvalbumin reporter mice.

**Results:** Juvenile *Pcdh19^+/-^* females lacked spontaneous seizures but displayed lower seizure thresholds and more severe seizures during hyperthermia. Seizure susceptibility did not differ from controls after flurothyl exposure. *Pcdh19^+/-^* females also exhibited segregation of GFP+ cells in the cortex, hippocampal CA1 region and medial ganglionic eminence, with a marked reduction of parvalbumin-positive interneurons in the CA1 hippocampal region. Although parvalbumin interneuron density was unchanged in the *Pcdh19^+/-^* female cortex overall, localized decreases arose in GFP- (*Pcdh19* knockout) cortical stripes.

**Interpretation:** Juvenile PCE mice exhibit seizure susceptibility to hyperthermia and disrupted the distribution of parvalbumin-expressing interneurons in the hippocampus and cortex. These findings suggest focal parvalbumin interneuron alterations may contribute to PCE pathophysiology.

## Introduction

PCDH19-clustering epilepsy (PCE) is a severe developmental and epileptic encephalopathy and one of the most common monogenic epilepsies^1^. PCE is characterized by cognitive impairment and intractable seizure clusters, often provoked by febrile illness, with onset in the first few years of life and an increased risk of neuropsychiatric disorders including schizophrenia in adulthood^2,3^. *PCDH19* is an X-linked gene that encodes a transmembrane cell adhesion molecule, critical for cell-cell interactions during brain development^4^. PCE affects females and rare mosaic males, while hemizygous males expressing only mutant *PCDH19* do not develop epilepsy, though neuropsychiatric features have been reported^5^. A leading hypothesis to explain this unique inheritance pattern is “cellular interference” associated with random X-inactivation (or somatic mutations) in which cells expressing only wild type (WT) and those expressing only mutant *PCDH19* fail to interact properly during brain development to cause disease^6^. This phenomenon is found in rodent and human induced pluripotent stem cell (iPSC) PCE models in which cells lacking *Pcdh19* tend to cluster together, as do WT cells^7–10^. However, the significance of this clustering pattern in the development of the different clinical manifestations of PCE remains unclear.

Loss of Pcdh19 in cultured neuronal or *in vivo* mouse models decreases synaptic connections and firing of excitatory neurons^11,12^. Decreased excitation alone is not consistent with a seizure phenotype, however, raising the possibility that altered excitatory/inhibitory balance is needed for full expression of the PCE phenotype. Seizures, in particular, may be better explained by deficits in cortical interneurons, a hypothesis that we sought to explore with this model.

PCDH19 also influences neuronal signal transduction as the cytoplasmic region of the protein binds to GABAAR-alpha subunits to regulate receptor surface availability, suggesting a role in GABAAR intracellular trafficking^13^. Electrophysiological studies of cultured hippocampal neurons have found that *Pcdh19* knockdown reduces tonic inhibitory currents^14^. Excitatory synapses and neurite branching are altered in *Pcdh19* mosaic neuron populations *in vitro*, further suggesting a role in synaptogenesis, likely through cell-cell adhesive interactions^12^. *Pcdh19* heterozygous (HET) female mice show presynaptic dysfunction of dentate granule cell-CA3 synapses that likely arise from mismatched interactions between pre- and postsynaptic *Pcdh19*/N-cadherin complexes. These mismatches are caused by *Pcdh19* mosaicism, and the disrupted synaptic cell-cell interactions alter ß-catenin signaling to reduce vesicular glutamate release^7^.

In addition to the function of GABAergic signaling in adult brain neurotransmission, GABA also plays a critical role in brain development, as migration and morphologic maturation of neurons rely on the trophic depolarizing action of GABAARs^15^. Several PCE models have shown subtle alterations in GABAergic neuron distribution within the brain ^16–18^. Despite recent advances in understanding the varied roles of *PCDH19*, however, the mechanisms by which mosaic expression leads to the phenotypic spectrum of PCE remains unclear. This study sought to utilize a novel mouse model of PCE to further evaluate the unique cell-clustering phenomenon of PCE and how it may relate to seizure susceptibility and alterations in cortical interneuron development.

## Methods

### Animals

All animal procedures were performed following protocols approved by the Institutional Animal Care and Use Committee of the University of Michigan and in accordance with the U.S. Public Health Service’s Policy on Humane Care and Use of Laboratory Animals. Mice were maintained using a constant 12 h light/dark cycle with access to food and water *ad libitum*. *Pcdh19*-null mice^7^ were maintained on a C57Bl6/J background (Jackson lab). Female *Pcdh19^-/-^* or *Pcdh19^+/-^*mice were bred with X-GFP male mice^19^ (Jackson lab #003116) to generate the female *Pcdh19^+/-^* PCE model, *Pcdh19^-/y^* males and *Pcdh19* wild-type control littermates (Fig. 1a). WT C57Bl/6J female mice were also bred with X-GFP males to produce control females. For conditional transgenic reporter studies, parvalbumin (PV)-cre mice (B6.129P2-*Pvalb^tm1(cre)Arbr^*/J, B6 PV^cre^, Jackson lab #017320)^20^ were bred into the *Pcdh19^-/-^* line and tdTomato (tdTom) cre reporter mice (B6.129S6-*Gt(ROSA)26Sor^tm14(CAG-tdTomato)Hze^*/J, Ai14, Jackson lab #007908)^21^ were bred into the X-GFP line, resulting in tdTom labeling of all PV-expressing neurons in offspring (Fig. 5A).

**Figure 1:**
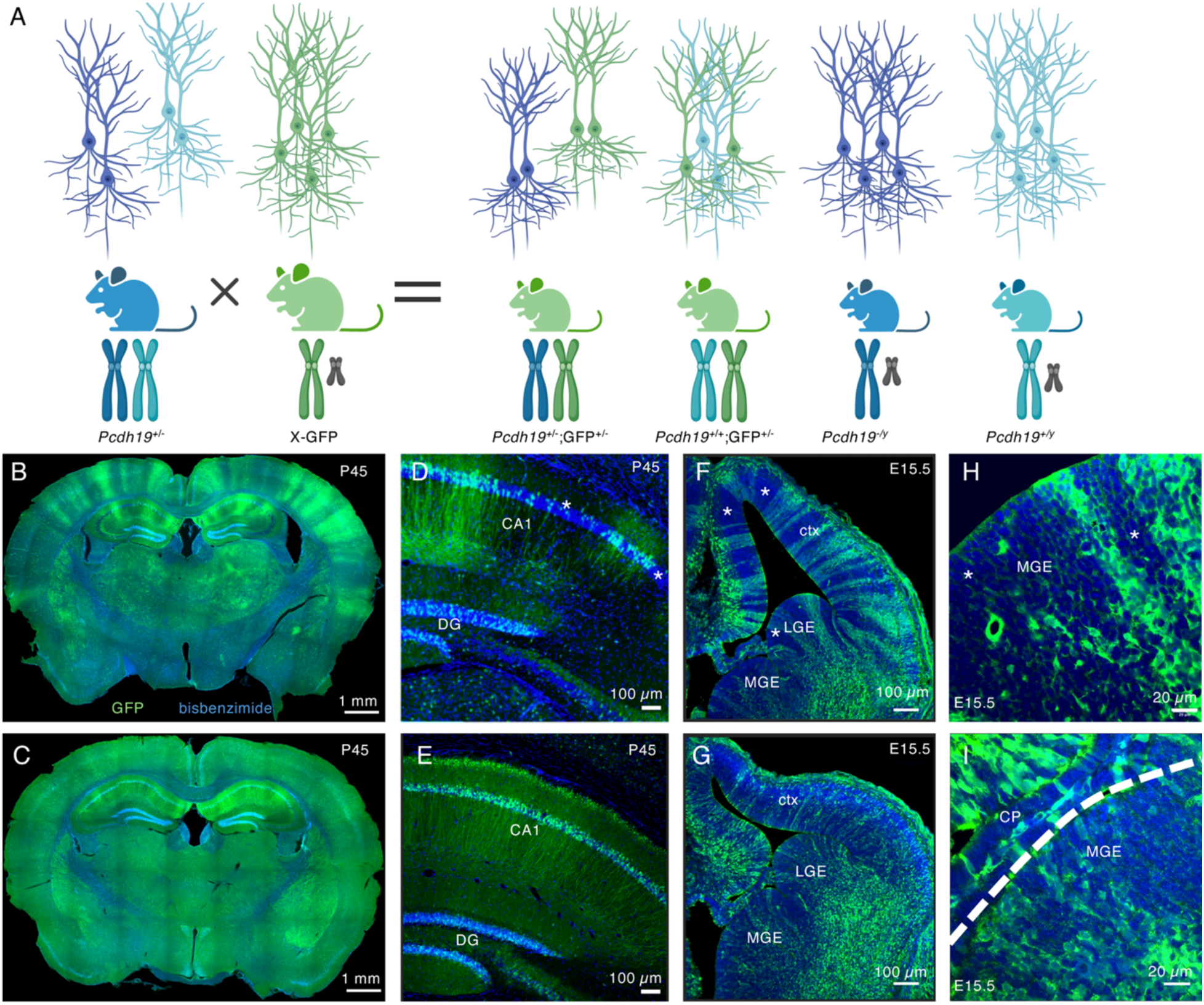
Breeding scheme and cell segregation in PCE mice. (A) *Pcdh19* heterozygous (HET) KO female mice (blue) with one KO *Pcdh19* allele (dark blue) and one WT allele (light blue) are crossed with an X-linked GFP male (green). Female offspring express a GFP on the paternally expressed X-chromosome, leading to mosaic GFP expression after random X-inactivation, and they are predicted to exhibit cell segregation (above). Male offspring are either *Pcdh19* hemizygous KO or WT, and neither are predicted to show cell segregation. (B-E) *Pcdh19* heterozygous female offspring (B) display a striking cell clustering pattern in the cortex and CA1 region of the hippocampus (D), while WT females lack this cell clustering phenotype (C, E). Male offspring do not express GFP (not shown). (F-I) The cell segregation pattern is seen early in development and by E15.5, there is a clear difference between HET (F) and WT (G) female mice, with clustering apparent in the cortex (ctx), lateral ganglionic eminence (LGE) and medial ganglionic eminence (MGE). High magnification images of the MGE in HET (H) and WT (I) female embryos at E15.5 further demonstrate the cell segregation pattern (the dashed line in (I) separates the choroid plexus (CP) and MGE). Examples of segregated regions with low-GFP (denoting silencing of the *Pcdh19* WT allele) in HET brains are marked by asterisks in D, F, and H. *Image created with Biorender.com*

### Thermal seizure induction

P15-16 mice (n=18-29 per genotype) were exposed to hyperthermia to induce seizures as per published methods^22^. Testing was digitally video recorded and stored on a secure server for offline analysis by a blinded observer. Seizures were scored on a modified Racine scale of 1-6 as follows: 1= staring, unresponsiveness, facial twitching; 2 = focal or clonic convulsion involving twitches or myoclonic jerk of a single limb, head nods, or backing; 3 = clonus of both forelimbs; 4 = rearing/uncontrolled hind limbs without loss of posture; 5 = loss of upright posture usually preceded by jumping/rearing; 6 = prolonged convulsion (> 30s of tonic/clonic convulsions with loss of posture) or death^22^. Grade 1 seizures were not scored, as they were difficult to confidently discern on video analysis. Mice were removed from the heat source and rapidly cooled on an ice pack after onset of a grade 5 or 6 seizure, or after 15 minutes with a body temperature of 42.5°C.

### Flurothyl seizure induction

Flurothyl seizure induction was performed similar to published methods^23^. Briefly, juvenile (P15-16) mice were individually placed in a sealed induction chamber. Flurothyl (2,2,2-trifluroethyl ether, Sigma-Aldrich) was pumped onto a gauze pad within the chamber at a constant rate of 20 µL/min. Latency to generalized tonic-clonic seizure was observed and events were video recorded for post-hoc review as necessary. Observers were blinded to genotype. Mice were immediately removed from the chamber following seizure induction and euthanized at the end of the session.

### Video/EEG recordings

Video/EEG recordings were performed similar to previously published methods^24^. Briefly, bilateral parietal screw electrodes were surgically implanted in juvenile (P21) HET, WT, and KO littermates (n=3-5 each). After a 7-day recovery period, simultaneous EEG recordings and infrared video monitoring were performed with a Natus recording system for 14 days. Seizures and interictal background were assessed manually by an experienced reader.

### Behavioral analysis

Adult (P90-150) litter matched (N=15 per group) mice were shipped to Scripps Research Institute and acclimated for 3 weeks prior to undergoing a battery of behavioral tests. Testing was designed to examine measures of anxiety, social interaction, sensory-gating, learning, and memory (see Supplementary Methods). Examiners were blinded to genotype at the time of testing. Genotype of 1 mouse could not be confirmed following testing, so the data were excluded for that subject. Statistical analysis was performed on all subjects and also for subgroups based upon sex with Student’s t-test and one-way ANOVA as appropriate to examine genotypic differences.

### Tissue processing and immunohistochemistry

Adult mice (P45-P90) were deeply anesthetized with pentobarbital and transcardially perfused with phosphate buffered saline followed by 4% paraformaldehyde (PFA). Brains were removed and post-fixed in 4% PFA overnight at 4°C and cryoprotected, frozen and cryosectioned at 40 µm thickness. Series of 5-6 sections, 240 mm apart, were processed as floating sections for immunofluorescence histochemistry (IHC) using the following primary antibodies: chicken anti-GFP (1:1000, Aves), mouse (Ms) anti-PV (1:500, Sigma), Ms anti-GAD67 (1:500, Millipore), Ms anti-somatostatin (SST; 1:200, Santa Cruz), and rabbit anti-GABA (1:500, Sigma). For PV-cre/TdTomato reporter visualization, rabbit anti-RFP primary antibody staining (1:500, Abcam) was used to enhance red fluorescence. Secondary antibodies (Alexa Fluor, 1:400 dilution, Invitrogen) used were: goat (Gt) anti-chicken 488, Gt anti-mouse 568, and Gt anti-rabbit 568. Nuclear counterstain was performed using bisbenzimide. For embryonic brain processing and IHC, embryos were dissected following maternal perfusion, fixed and cryprotected as above. Whole embryo heads were sectioned onto slides at 20 µm and stained with anti-GFP antibody. Nuclear counterstain was performed with bisbenzimide (Hoechst).

### Microscopy

IHC images of antibody-stained sections were acquired with a Nikon A1 confocal microscope under a 20x objective and 1.0x optical zoom with 2μm step size through the z-plane and the pinhole set at 1 Airy unit. Imaging of sections from transgenic reporter mice was obtained on a Nikon epifluorescence microscope. Whole brain sections were imaged at 10x and digitally stitched with Nikon NIS-Elements software.

### Image analysis

Hippocampal images were imported into Image J for quantification of interneuron cell density. The observer was blinded to experimental group (without the GFP channel). Cell density was calculated as the number of immunolabeled neurons per mm^2^ of the CA1 region. For each mouse, total number of cells on 3-6 sections were counted and divided by the total CA1 area examined. Cortex and hippocampal images from PV-cre/tdTom reporter lines were evaluated with custom-written masking scripts in Image J. Thresholds were set for analysis of “high-GFP” and “low-GFP” regions with automated cell counting of tdTom-positive neurons for PV or for cells immunostained for SST with far red-conjugated secondary antibody. The same script was employed to analyze overall cell density by counting bisbenzimide positive cells in each region. To analyze variability in masked regions, masks of 4 PCE mice were superimposed on anatomically similar sections of WT and knock-out mice for analysis of PV cell density.

### Statistical analysis

Statistical analyses for all experiments were performed using GraphPad Prism 10 software. Group means were compared by one-way ANOVA or Student’s t-test with significance set at p<0.05. Paired t-tests were used for within animal comparisons. For seizure susceptibility, survival curves were compared with a Log-rank (Mantel-Cox) test. Behavioral assays were analyzed with two-way ANOVA as appropriate.

## Results

### Cell segregation in the X-GFP PCE mouse model

Previously published PCE mouse models displayed a unique striping pattern of *Pcdh19*-positive (hereafter referred to as WT) and *Pcdh19*-negative (hereafter referred to as knockout, KO) neurons in the cortex of *Pcdh19*^+/-^ females due to random X-inactivation^7,10^. We therefore chose to utilize a commercially available X-linked GFP reporter mouse to label wild-type neurons after random X-inactivation of either the WT (GFP+) or KO (GFP-) X-chromosome (Fig. 1A). Indeed, in our model, brains from female PCE (*Pcdh19* HET) mice displayed a striking cell segregation pattern with GFP+/*Pcdh19*+ neurons segregated from GFP-/*Pcdh19*-neurons (Fig. 1B), which is not seen in WT female littermates (Fig. 1C). This pattern is seen as early as embryonic day (E) 10 (data not shown) and is maintained through adulthood. In the adult, the striping is clearly discernable in the cortex and the CA1 region of the hippocampus (Fig 1B-E). Interestingly, the CA3 region and dentate gyrus showed a random pattern of GFP+ and GFP-cells without segregation (Fig. 1D). Notably, consistent with the process of random X-inactivation, the cell segregation pattern within the cortex and hippocampal CA1 region varied between PCE mice, such that no 2 mice displayed the same pattern of GFP expression. In the embryo, cell segregation is evident in the cortex and ganglionic eminences (Fig. 1F-G), including within interneuron progenitor domains of the medial ganglionic eminence (MGE, Fig. 1H-I).

### Juvenile PCE mice are susceptible to hyperthermia induced seizures

Rodent models of PCE have failed to recapitulate spontaneous seizures, though they have been reported to have a lower seizure threshold in several assays^23^. We recorded simultaneous video/EEG in 5 awake behaving PCE mice and WT and KO littermates from P28-P42 and captured no spontaneous seizures or clear interictal abnormalities (data not shown). Patients with PCE typically present with intractable seizure clusters in the first few years of life, often provoked by fevers^4^. Given the clinical relevance of fever-provoked seizure clusters in young girls with PCE, we examined seizure thresholds with exposure to hyperthermia in juvenile mice. Hyperthermia exposure at P15-16 produced abnormal head jerks consistent with Racine Grade 2 seizures in the majority of mice tested by the end of the assay (98/102 mice), regardless of genotype. However, more severe seizures were induced in female PCE mice compared to male hemizygous KO or WT female and male controls (Fig. 2A). PCE female mice also displayed a lower temperature threshold for first head jerk than WT mice, but interestingly, KO males appeared to have an intermediate phenotype with a seizure threshold lower than WT males, but not significantly different from HET females (Fig. 2B). Survival curves of the 4 groups also showed that PCE (HET female) mice had the lowest seizure threshold, with KO males displaying an intermediate phenotype between the WT and HET females (Fig. 2C).

**Figure 2:**
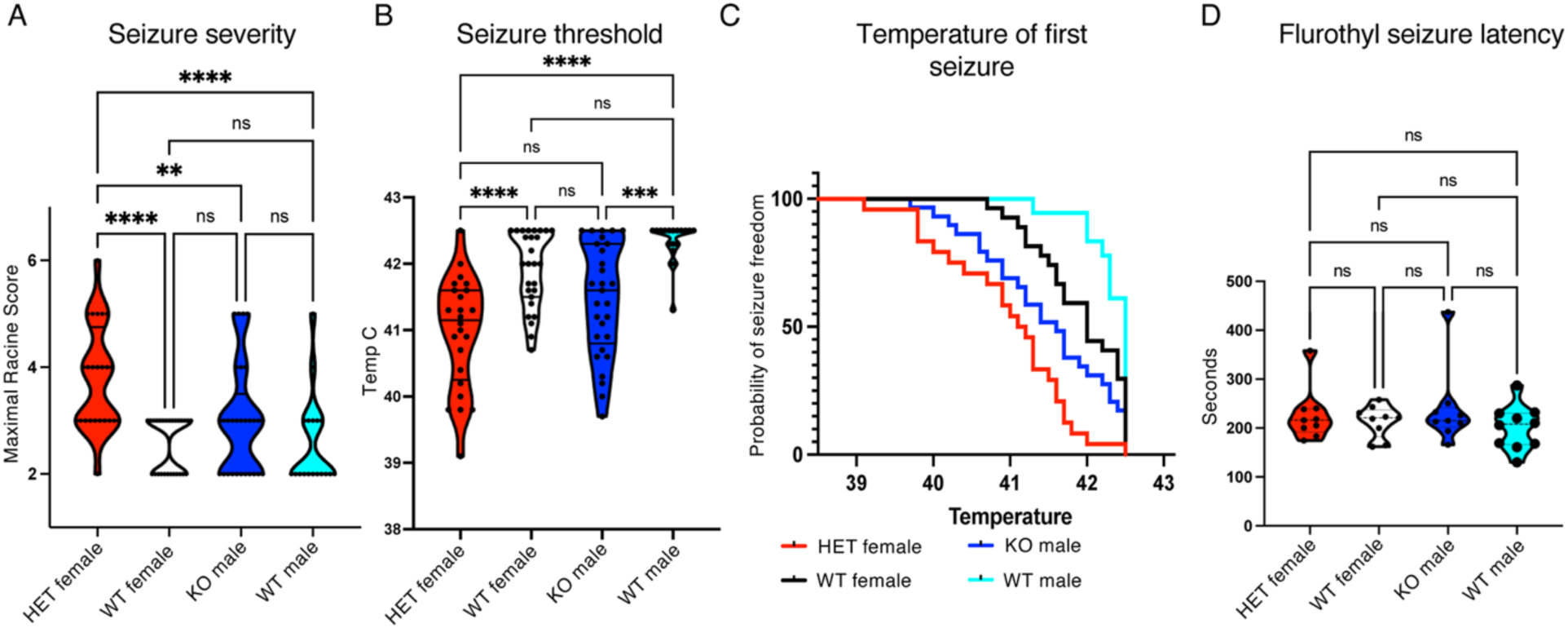
PCE mice exhibit more severe seizures and lower temperature thresholds with hyperthermia. (A) A hyperthermic seizure assay at P15-16 revealed more severe seizures in HET females compared to litter-matched controls as scored on a modified Racine scale (one-way ANOVA). (B) HET females also had a lowered seizure threshold compared to both WT controls, and the hemizygous male KO group displayed a significantly lowered seizure threshold compared to WT males (one-way ANOVA). (C) Survival curve of temperatures that induced the first seizure show significantly lower temperatures in HET females compared to WT controls, while KO males displayed an intermediated threshold curve (Logrank (Mantel-Cox) test, HET female vs. WT female, p<0.0001, KO male vs. WT male, p=0.0004, HET female vs. KO male, p=0.0236). (D) Flurothyl exposure in mice at P15-16 did not reveal any significant differences between groups for seizure latency (one-way ANOVA). (A-C, N=24, 27, 29, 18. D, N=9 per group). n.s – p>0.05, **p<0.01, ***p<0.001, ****p<0.0001.

Other mouse models of genetic epilepsies with a strong predilection for febrile seizures, such as genetic epilepsy with febrile seizures plus (GEFS+), have shown a seizure sensitivity to hyperthermia in juvenile cohorts that is not found in adults^22^. Given our findings of hyperthermia sensitivity in juveniles and previous reports of a sensitivity to fluorothyl-induced seizures in adult PCE mice^23^, we examined fluorothyl sensitivity at the juvenile (P15-16) timepoint. This testing revealed no difference in seizure susceptibility between the 4 groups, albeit with greater variability in the HET female and hemizygous male KO mice compared to WT littermates (Fig. 2D).

### PCE mice do not have deficits in learning or memory

Published studies of behavioral alterations in PCE animal models revealed variable deficits that are not consistent between models^7,11,25–27^. We additionally performed a battery of tests in adult mice to specifically assay for behavioral disorders similar to those observed in patients with PCE who variably display cognitive abnormalities, autistic features and an increased risk of schizophrenia development in adolescence^28–30^. Assays to assess anxiety-like or repetitive behaviors including elevated plus maze, open field, and marble burying tests revealed no differences between WT and PCE female mice (Fig S1A-C), whereas KO male mice buried significantly fewer marbles than their PCE and WT littermates (Fig. 3A) and WT male mice spent more time in the center of the elevated plus maze than both WT and PCE females (Fig S1A). A test of social interaction showed no differences between groups (Fig. 3B). Assessments of learning and memory including novel object recognition, Morris water maze, and conditioned fear testing also revealed no differences among the genotypes (Fig. 3C, S1D). Finally, analysis of acoustic startle/pre-pulse inhibition (PPI) testing showed subtly significant differences between genotypes, with several increased amplitudes of startle responses in HET female mice compared to their wild-type female littermates, but no significant difference in PPI (Fig 3D, S1E). There was no significant difference in responses of KO males compared to wild-type male littermates.

**Figure 3:**
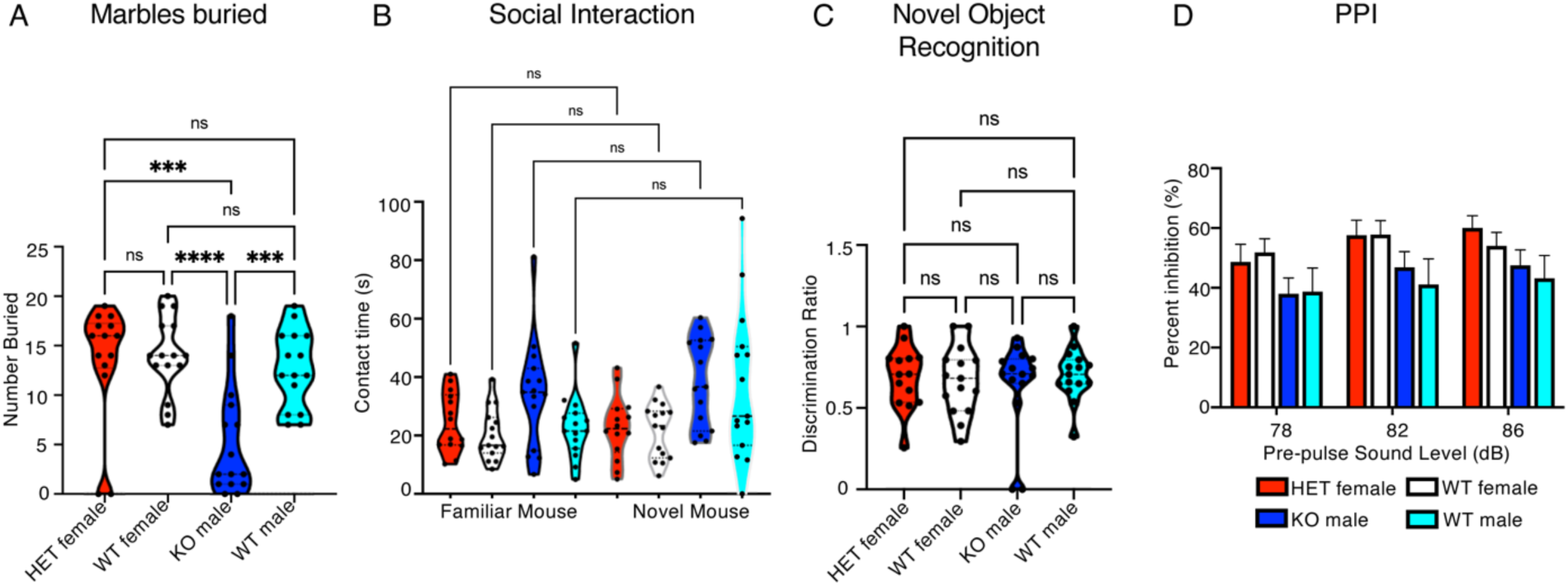
A battery of behavioral testing revealed few differences between genotypes. (A) Pcdh19 KO males buried significantly fewer marbles during the task than litter-matched HET female, WT female, and WT male mice (one-way ANOVA), but there was no significant difference between HET and WT females (ns – p>0.05, ***p<0.001, ****p<0.0001). (B) Social interaction testing showed no difference in the amount of time spent in contact with a familiar versus novel mouse across any genotypes (one-way ANOVA). (C) Novel object recognition was similar between genotypes and sexes (one-way ANOVA) as was pre-pulse inhibition (PPI) testing (D, p>0.05 two-way ANOVA).

### PV interneuron density is altered in the hippocampus and cortex of PCE mice

To explore potential mechanisms for the increased seizure susceptibility in PCE mice, we next assayed for alterations in interneurons. A previously published PCE mouse model found only subtle changes in interneuron density in the cortex of PCE mice^16^. We therefore initially chose to examine the hippocampus, as imaging studies of patients with PCE suggest that subtle anatomical differences are present in limbic structures compared to controls^31^ and the robust cell segregation we find in the CA1 region raises the possibility that *Pcdh19* mosaicism alters hippocampal development. In addition, though multiple seizure types have been described in PCE, many patients exclusively experience temporal lobe seizures^32^. We therefore performed immunohistochemistry in adult *Pcdh19* HET and WT female mice to examine cellular density of the main MGE-derived interneuron subtypes with PV and SST immunolabeling, as well as total interneuron numbers by immunostaining for GAD67 and GABA (Fig. 4A). Given the clinical phenotype in heterozygous females, we limited this initial evaluation to females only. Evaluation of interneuron density in the CA1 region of the hippocampus, an area with robust cell segregation, revealed significantly fewer PV-positive cells in PCE mice when compared to WT females (Fig. 4B). No significant differences were found in cell density of the other main MGE-derived interneuron subtype, SST, or for total interneurons immunolabeled with GAD67 or GABA. To further examine PV interneurons in CA1, we crossed B6 PV^cre^/Ai14 transgenic reporter mice into our PCE mouse model (Fig. 5A-B). Consistent with our IHC results, PCE mice exhibited a lower density of tdTomato-positive cells in the CA1 region compared to WT and KO littermates (Fig 5C-D). Consistent with previous work using a different PCE mouse model^16^, we found no significant change in PV interneuron density in the cortex (Fig. 5E). Together, these findings suggest that the lower susceptibility to hyperthermia-induced seizures in PCE mice may be associated with a decrease in the density of PV interneurons in the hippocampal CA1 region, but not in the cortex.

**Figure 4:**
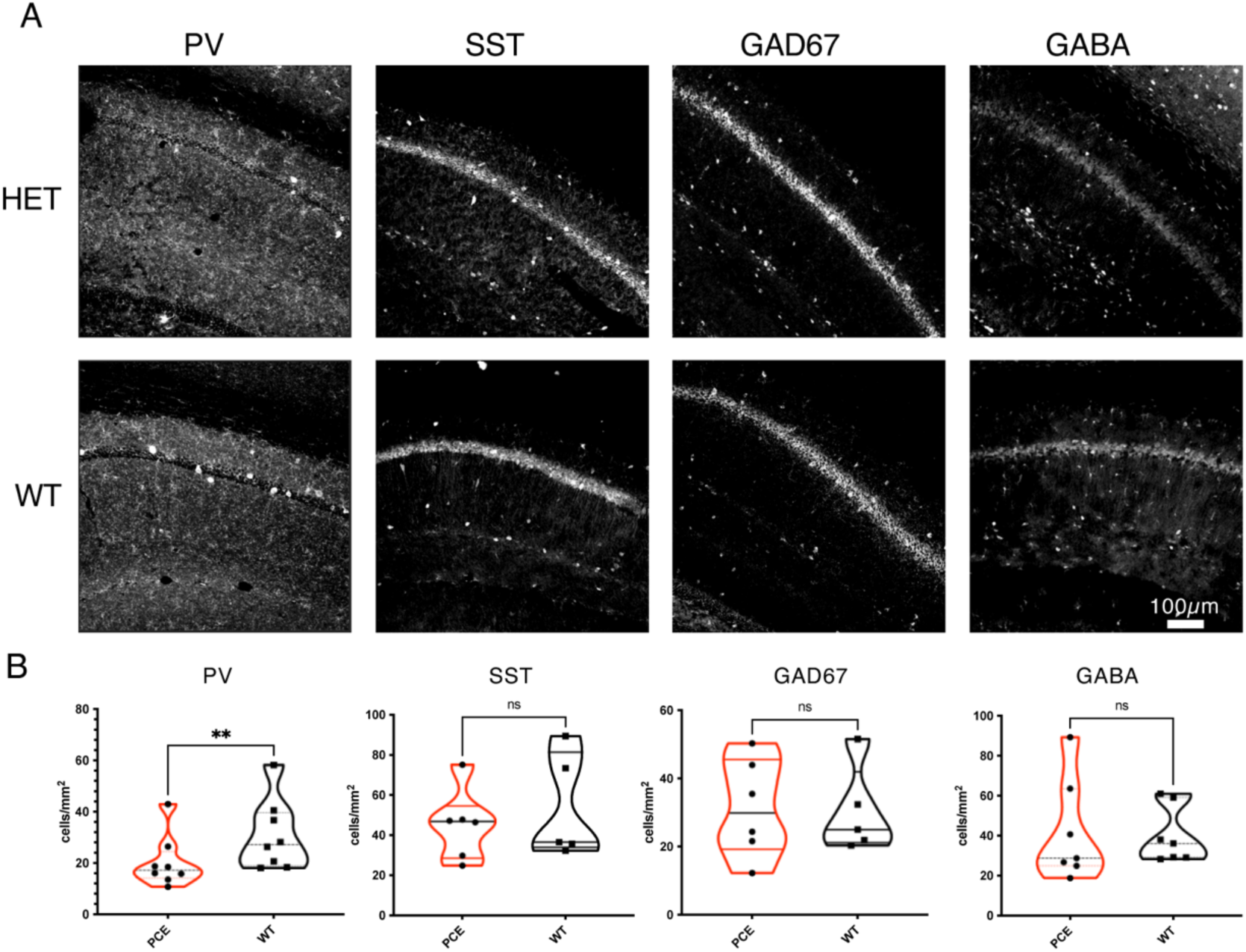
Interneuron density in the hippocampal CA1 region of adult PCE (top) and WT (bottom) mice. (A) Immunolabeling for parvalbumin (PV), somatostatin (SST), GAD67, and GABA in *Pcdh19* HET (top) and WT (bottom) female mice. (B) Quantification of interneuron density revealed a significantly lower density of PV neurons in PCE mice compared to WT (**p<0.01), with no significant differences in SST (p=0.55), GAD67 (p=0.90), or GABA (p=0.88). N=6-8 mice per group. Unpaired t-test.

**Figure 5.**
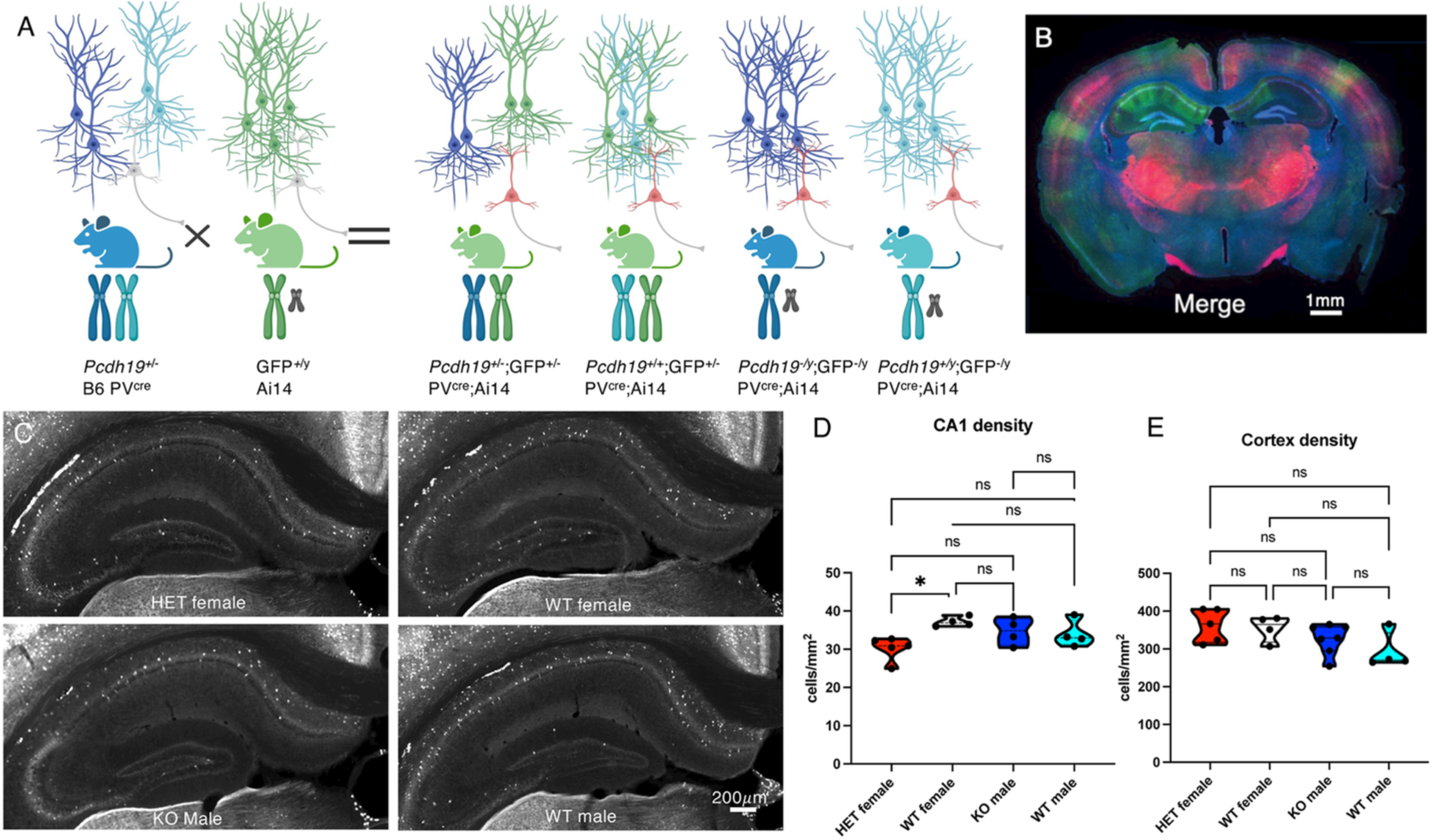
(A) B6 Pv^cre^ and Ai14 transgenic reporter mice were bred into our PCE model, resulting in expression of TdTomato on PV expressing interneurons (B). Quantification of PV density in the CA1 region of the hippocampus (C) was consistent with our IHC findings, with HET females displaying a lower density of PV reporter-expressing interneurons than WT female littermates (D - * p<0.05, n.s. p>0.05, one-way ANOVA). (E) Quantification of PV density in the cortex was not significantly different between groups (n.s. p>0.05, one-way ANOVA). *Image created with Biorender.com*.

Notably, our PCE model enables the evaluation of interneuron distribution in the context of *Pcdh19* expression by examining PV cortical interneuron distribution in relation to GFP labeling. Within PCE females, we utilized a custom masking script to evaluate PV density in highly GFP-expressing stripes (as a surrogate for WT *Pcdh19* expression) versus predominantly unlabeled (KO) stripes in an unbiased fashion using the PV reporter line (Fig. 6A-B). We first examined the hippocampal CA1 region and found that PV interneuron density in high-GFP versus low-GFP regions was not significant different (Fig. 6C). We then applied the same masking protocol to the cortex (Fig. 6D-E) and observed a significant alteration in the density of PV positive interneurons in the high-GFP versus low-GFP regions of the PCE cortex (Fig. 6F), such that there was a significantly lower density of PV interneurons in *Pcdh19*-negative (low-GFP) regions when compared to *Pcdh19*-positive (high-GFP) regions (Fig. 6G). Overall cell density, as examined by bisbenzimide labeling, was not different between the high-GFP and low-GFP regions (Fig. 6H), but the percentage of PV positive cells was significantly greater in the high-GFP regions (Fig. 6I).

**Figure 6:**
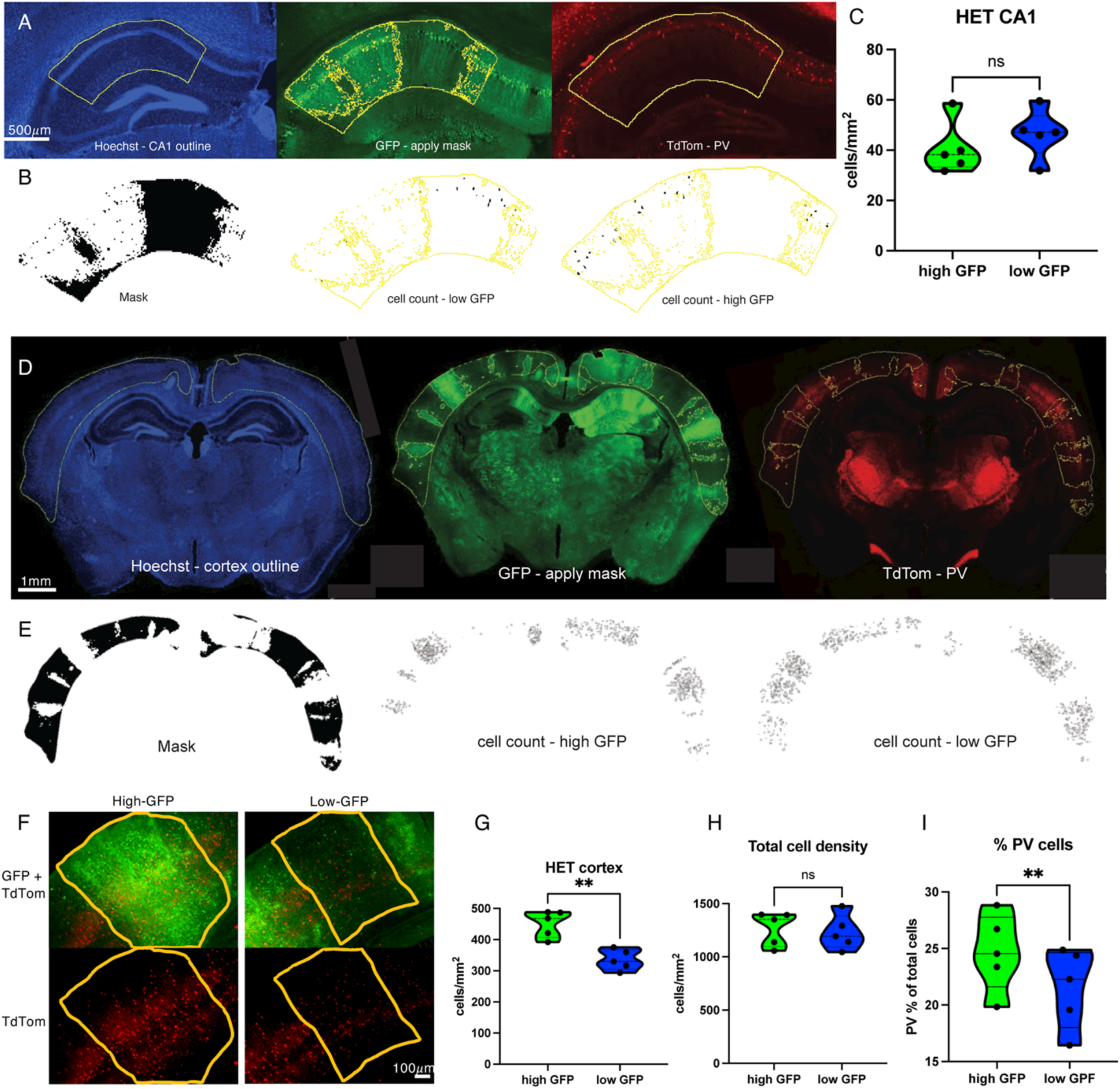
Parvalbumin interneuron density is decreased in *Pcdh19* KO cortical, but not hippocampal, stripes. (A) In *Pcdh19* HET females expressing the PV-cre; tdTomato reporter, segmentation masks were created to visualize PV interneurons in high-GFP versus low-GFP expressing regions of hippocampal CA1. (B, C) Automated cell counting revealed no difference in PV density in the CA1 region between high-GFP and low-GFP expressing regions (ns, p=0.1427, paired t-test). (D-F) The same segmentation mask was employed in the cortex to examine PV density in high-GFP versus low-GFP cortical stripes. (G) Quantification revealed a significantly lower PV reporter-labeled cell density in low-GFP regions of the cortex compared to high-GFP regions (**, p<0.01, paired t-test). (H) A similar quantification of total cell densities in the high GFP versus low GFP regions was showed no significant difference (ns, p>0.05, paired t-test). (I) The percentage of cells that were TdTomato-positive was higher in the high GFP regions versus the low GFP regions (**, p<0.01, paired t-test). N=5 mice.

To examine if this finding was simply a result of regional differences in PV-density, we superimposed the GFP masks of PCE mice onto anatomically equivalent sections from WT female and WT and hemizygous KO male mice to create a “false mask” (Fig. S2A-B). This evaluation revealed no significant difference in the PV density between the masked regions when superimposed onto control brains (Fig. S2C). Furthermore, we examined the within animal ratio of PV interneurons in the high-GFP:low-GFP regions in PCE mice and in hemizygous males and WT females and males, using the “false masks” for the latter 3 groups. This assessment also confirmed that the alteration in PV interneurons was specific to PCE mice and their associated *Pcdh19* mosaicism (Fig S2D). SST immunostaining was then performed on sections from the same B6 PV^cre^/Ai14 PCE mice from a single litter to assess for *Pcdh19* mosaicism-associated changes in the other major MGE-derived interneuron subtype. Overall SST interneuron density in the cortex of PCE mice was not different from litter-matched controls (Fig. S3A-B). Employment of the same custom masking script with the SST channel also revealed no difference in SST density between the high-GFP and low-GFP regions of the cortex (Fig. S3C). These findings suggest that mosaic expression of *Pcdh19* selectively reduces PV interneuron density in KO cortical regions, potentially leading to multifocal areas of cortical hyperexcitability.

## Discussion

We describe a novel model of PCE that involves crossing *Pcdh19* KO mice with a transgenic X-linked GFP reporter line, the latter serving as a surrogate marker of random X-inactivation to identify brain regions with WT versus mutant *Pcdh19* expression. We used this model to examine PCE cell segregation patterns and to explore seizure susceptibility and other behavioral phenotypes. To better understand potential causes of cortical hyperexcitability in PCE, we also measured interneuron density in cortex and hippocampus. Our model displayed robust segregation of the cell population expressing WT *Pcdh19* (GFP+) from that lacking *Pcdh19* (unlabeled KO) in cortex, hippocampal CA1 and ganglionic eminences. The cell segregation arose during embryonic development and persisted into adulthood. PCE mice also displayed a lower threshold and increased severity of hyperthermia-induced seizures without consistent abnormalities in learning and memory or other cognitive domains. Notably, we found decreased PV interneuron density in the CA1 region of the hippocampus, irrespective of GFP labeling, and within *Pcdh19* KO (GFP-negative) cortical stripes. These findings suggest a novel epilepsy mechanism with focal regions of impaired inhibition that may lead to seizures and developmental deficits in PCE.

Similar to previous reports^10,33^, the cell segregation phenotype in our model is visible as early as E10, around the onset of *Pcdh19* expression^34^. Due to random X-inactivation, each mouse displayed different patterns of GFP expression, underscoring the inherent variability in PCE mouse models and potentially underlying the variable phenotypic expressions in patients with PCE^2^. Interestingly, we did not observe the segregation pattern in the CA3 region or dentate gyrus of the hippocampal formation, despite evidence that *Pcdh19* is expressed in these regions^7^. The mechanism by which *Pcdh19* mosaicism leads to cell segregation is unknown, but likely involves the early expression of PCDH19 in ventricular zone radial glia^8,10,35^ and its role in cell adhesion and neuronal migration. The lack of segregation in specific regions of the mouse brain may reflect temporal variability in *Pcdh19* expression in differing cell populations or, for the dentate gyrus, may relate to the fact that dentate granule cells migrate tangentially to form the granule cell layer rather than as radial units^36^. Our model provides potential advantages over previously reported PCE models^7,10,16,17^ for studying the role of *Pcdh19* in neuronal development and synaptic transmission given that it enables visualization of mosaic cell populations in live tissue.

We show that PCE mice have a lower seizure-threshold when exposed to hyperthermia, consistent with patient susceptibility to fever-induced seizures. Behavioral phenotyping in PCE models has been challenging, as many mouse models have displayed only subtle phenotypes that vary between models^7,11,23,26^. Our PCE mouse model did not develop spontaneous seizures, similar to other mouse models of PCE and other genetic epilepsies^10,23,37^. However, the susceptibility to hyperthemic seizures in juvenile mice may be a clinically relevant biomarker, given the high prevalence of fever-provoked seizures in patients with PCE^38^. A juvenile, age-related susceptibility to hyperthermic seizures has been observed in other mouse models of genetic epilepsy^22^ and may represent a developmentally important window. Notably, our *Pcdh19* female HET and hemizygous male KO mice both displayed a significant amount of variability in seizure threshold and severity compared to WT controls. In HET mice, this effect may reflect the wide variability in *Pcdh19* expression patterns, potentially causing more significant variability in local circuits. The “intermediate” phenotype seen in our male KO mice is similar to other studies ^23,25,33^ and suggests that complete *Pcdh19* loss of function may also alter neural circuit excitability, though likely to a lesser degree than in female patients who are heterozygous or males with somatic mosaicism. Evaluation of seizure thresholds with flurothyl exposure did not reveal any genotypic differences in our cohort of mice, contradictory to a previously published study^23^. Additional behavioral phenotyping of our adult PCE mice also showed minimal differences in a battery of behavioral tests. Previous behavioral testing of PCE models has shown inconsistent deficits^7,16,26,33^. The differences in findings between studies in PCE mice may be age-related, the result of variable mosaicism from random X-inactivation patterns in each cohort of mice, or related to the genetic background of the various mouse models, as background strain is known to play a significant role in seizure susceptibility in epilepsy models^39,40^.

To investigate potential mechanisms underlying hyperthermia-induced seizure susceptibility, we examined interneuron density or distribution in PCE mice, with a focus on brain regions with observed cell segregation. Physiological studies of *PCDH19* mosaicism indicate that it disrupts synaptic contacts between *Pcdh19* knockout and wildtype neurons with overall reduced excitatory neuronal transmission^7,11^. These abnormalities are unlikely to result in seizures. Inhibitory networks are tightly controlled, such that reduced neuronal transmission from PV cortical interneurons may lead to altered local circuits and result in various neurologic manifestations. The decreased PV interneuron density within the hippocampus and KO cortical stripes may thus lead to the increased susceptibility to hyperthermia-induced seizures. Previous work examining interneurons in mouse models of PCE has shown subtle changes in the cortical laminar distribution of PV neurons in the cortex^16^, but interneuron alterations in the hippocampus have not been previously explored. Given the clinical relevance of limbic seizures in patients with PCE and imaging findings suggesting volumetric reduction in the hippocampal subfields^31,32,41^, in addition to the striking cell clustering pattern in the CA1 region that we observed, we initially focused on GABAergic interneuron densities in this region. Though we did not find any differences in total interneuron density in the PCE mice versus WT, it was intriguing to find a lower density of PV-positive interneurons in the PCE mice, suggesting that PV-specific interneuron dysfunction may play a role in PCE pathophysiology. Notably, PCDH19 is expressed in doublecortin-enriched nests (DENS) that are present in human, but not mouse, MGE^42^.

PV-expressing GABAergic interneurons play a vital role in feedforward and feedback synaptic inhibition, and alterations have been implicated in numerous genetic and acquired epilepsies^43,44^. PV cells in our model were less densely distributed in the hippocampus of PCE mice, while maintaining the same density in the cortex when compared to WT controls. This finding led us to pursue a deeper analysis of the PV interneuron distribution in the cortex in the context of the GFP striping pattern. Overall PV density in the cortex of PCE mice did not differ from controls. However, when further examining PV interneuron density in the context of GFP expression, cortical stripes with low GFP expression, denoting largely *Pcdh19* KO regions, displayed a lower PV interneuron density than WT cortical stripes with high GFP expression. This finding raises the possibility that seizures in PCE arise from focal regions of cortical disinhibition, a novel potential epileptogenic mechanism related to genetic mosaicism. Prior studies have shown that *Pcdh19* mosaicism leads to altered interneuron distribution and migration in zebrafish and MGE explant models^17,18^. The focal decrease of PV cortical interneurons in KO regions may reflect an alteration of precursor proliferation, migration, cell fate specification or survival. Moreover, we do not yet know whether the differences involve all PV interneuron subtypes (e.g. basket, axo-axonic and translaminar) or are biased to a specific subtype or cortical layer. Imaging of the embryonic mouse brain reveals neuronal segregation in both embryonic excitatory neurons and early-born inhibitory neurons prior to tangential migration. This suggests that “cellular interference” could play a role in interneuron development and migration, leading to decreased inhibitory neuron density in the juvenile mouse hippocampus and a hyperexcitability phenotype. Alternatively, loss of *Pcdh19* has been shown to decrease excitatory synaptic density^12^. It is possible that focal cortical hypoexcitability in *Pcdh19* KO regions therefore results in a compensatory decrease in PV-interneuron density through synaptic pruning and apoptosis.

This work establishes a novel model of PCE with the ability to visualize regions of *Pcdh19* expression in both live and fixed tissues. The focal cortical alterations of PV interneuron density that we identified may represent a novel epilepsy mechanism resulting from mosaicism. These alterations may also contribute to the variable PCE phenotypes observed in clinical practice. Our model can serve as a platform for further work to understand the contribution of PV-interneurons to PCE phenotypes, how these alterations arise in development, and for the identification of novel targets to treat PCE.

## Acknowledgements

Grant support for this study from NINDS, K08NS124937 (JZ), the American Epilepsy Society/PCDH19 Alliance (JZ), and the University of Michigan Department of Pediatrics via the Jeanette Farrantino Investigator award and Elizabeth E. Kennedy Children’s Research Award. The authors would like to thank Judy Huyhn, Zeeshan Bhalwani, Zoe Srackangast, Carla Rivera Pacheco, and Emily Langlois for technical assistance with these studies. Behavioral testing was performed at Scripps Research Institute with thanks to Dr. Amanda Roberts. Appreciation to Dr. Lori Isom, Dr. Chunling Chen, and Dr. Sundeep Kalantry for assistance with transgenic mice. Thank you to Dr. Louis Dang for critical review of the manuscript.

## Author Contributions

JZ: Conceptualization and design, data acquisition and analysis, manuscript drafting. JH: Data acquisition and analysis; manuscript editing. ND: Data acquisition and analysist, manuscript editing. EM: data acquisition, analysis and manuscript editing. KJ: Data acquisition and analysis, manuscript editing. HU: Experimental design, manuscript editing. JMP: Conceptualization and design, manuscript writing and editing.

## Data Availability

All data are available upon request.

