## Supplemental materials for "Hyperthermic Seizure Susceptibility and Focal Decreases in Parvalbumin-Expressing Cortical Interneurons in a Mouse Model of PCDH19-Clustering Epilepsy"

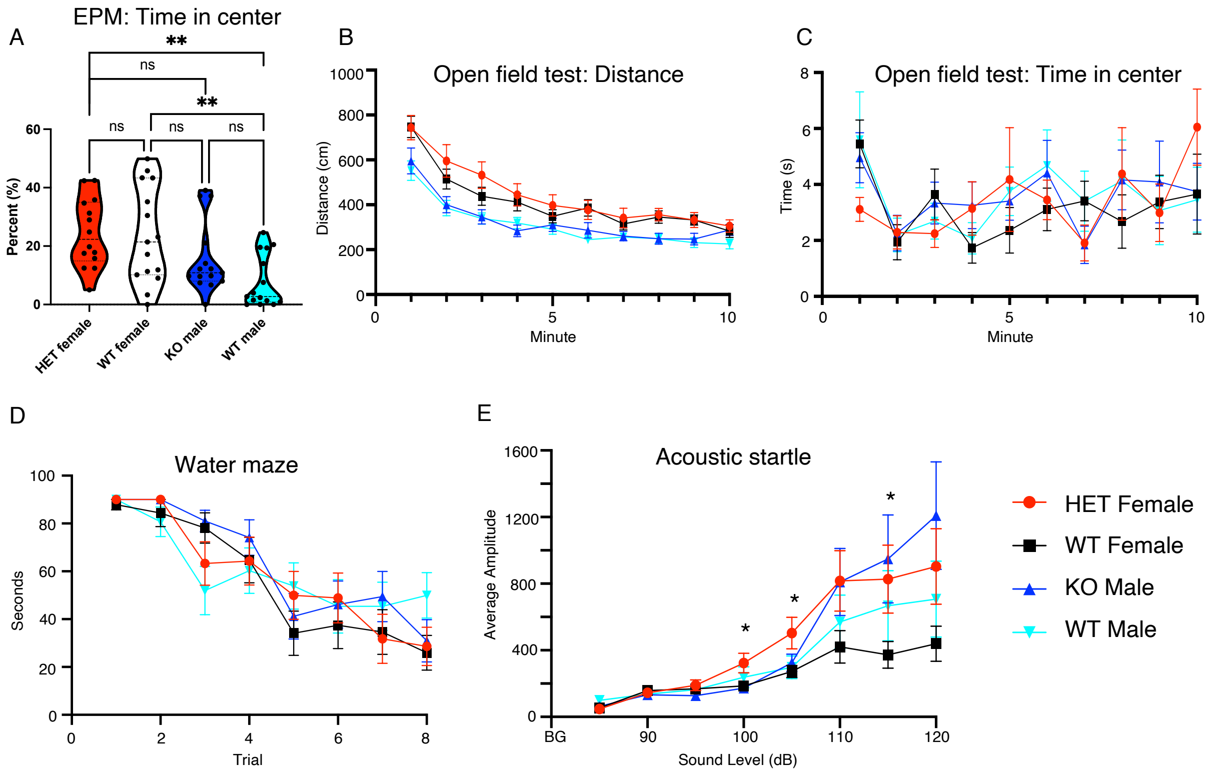


Supplemental Figure 1: (A) Elevated plus maze (EPM) testing revealed no differences in time in center between genotypes, though there was a sex difference with WT males spending significantly less time in the center than females (**$p<0.01$, ns – $p>0.05$, one-way ANOVA). Open field testing showed no differences in distance traveled (B) or time in center (C) between genotypes ($p>0.05$, two-way ANOVA). Water maze testing revealed no difference in time to reach the platform between genotypes (D, $p>0.05$, two-way ANOVA). Acoustic startle testing (E) showed significantly higher amplitude startle between HET females and WT females at several sound levels (*$p<0.05$), with no significant difference between KO and WT males ($p>0.05$, two-way ANOVA).

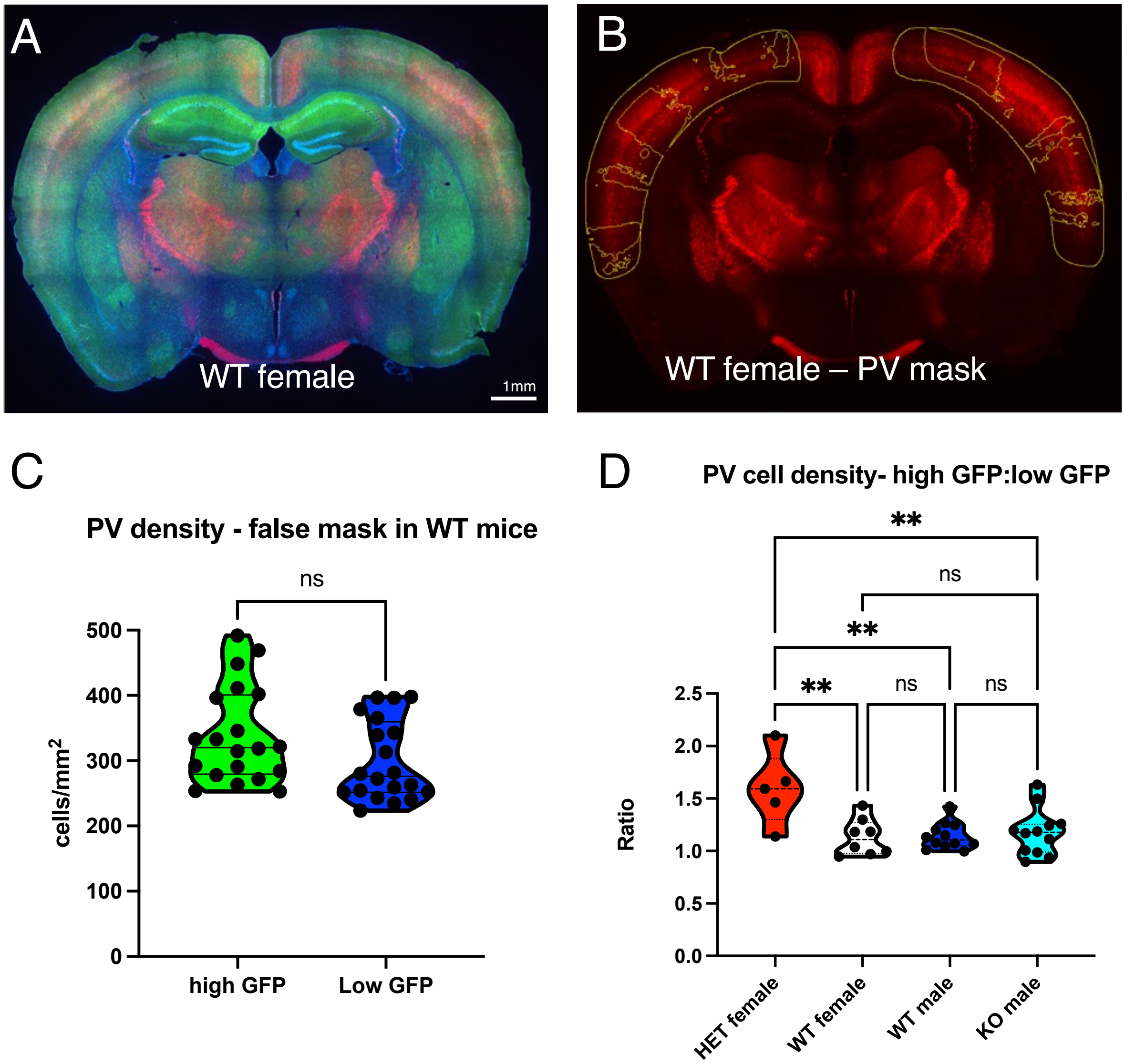


Supplemental Figure 2: (A-B)To assess if PV density alterations discovered in *Pcdh19* HET females were due to normal anatomic variation, we applied the GFP masks from HET females to similar anatomic sections of WT and KO control sections. (C) Direct comparison of tdTomato-labeled cell density using the false masks in WT mice revealed no significant difference in the high-GFP mask vs. low-GFP mask regions (ns, $p>0.05$, paired t-test). (D) The ratios of PV reporter-labeled cell density in high-GFP:low-GFP mask regions was determined for each individual mouse. This analysis revealed a significantly higher ratio in the HET females when compared to KO and WT litter mates (**, $p<0.01$; ns, $p>0.05$; one way ANOVA).

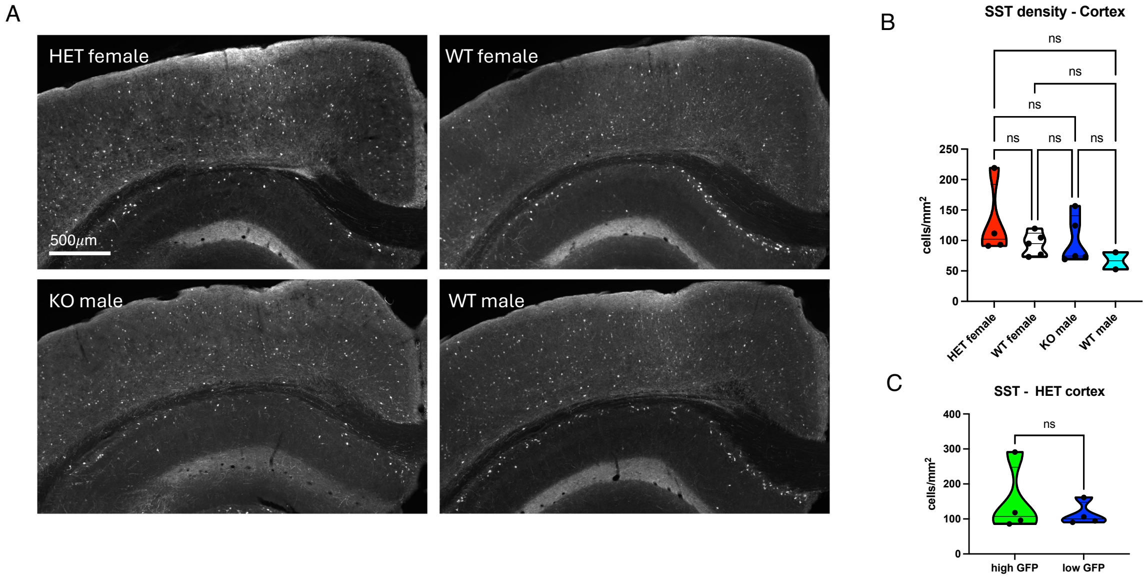


Supplemental Figure 3: (A) Immunostaining for SST was performed and analyzed with the same high GFP/low GFP masking script utilized for PV density analysis. (B) There was no difference in SST density in the cortex between the four groups of mice (ns, p>0.05, one-way ANOVA). (C) In heterozygous females, there also was no difference in SST density associated with cortical stripes having high versus low GFP expression. (ns, p>0.05, paired t-test).

**Supplementary methods**

Elevated plus maze test. Methods were modified from (1). Mice were placed in the center of the testing apparatus with four arms (5×30 cm) at right angles to each other, elevated 30 cm from the floor. Two arms had 16-cm black plastic walls (closed arms), and two arms had 16-cm clear plastic walls (open arms). Behavior was video recorded for 5 min and time spent in open arms was recorded.

Open field test. The apparatus was a square white Plexiglas (50 x 50 cm) open field illuminated to 600 lux in the center. Each animal was placed in the center of the field and several behavioral parameters (distance traveled, velocity, center time, frequency in center) were recorded during a 5-minute observation period and analyzed using Noldus Ethovision XT software.

Social interaction test. The social interaction apparatus is a rectangular, three chambered Plexiglas box, with each chamber measuring 20 cm x 40.5 cm x 22 cm (L x W x H). Dividing walls are clear with small semicircular openings (3.5 cm radius) allowing access into each chamber. The middle chamber is empty, and the two outer chambers contain small, round wire cages (Galaxy Cup, Spectrum Diversified Designs, Inc., Streetsboro, OH) during testing. Mice were habituated to the entire apparatus with the round wire cages removed for 5 min. To assess sociability, mice were returned to the middle chamber, this time with a stranger mouse (same-sex, age-matched C57BL/6J, habituated to the wire cage) in one of the wire cages in an outer compartment and another identical wire cage in the opposite compartment. Time spent in the chamber with the stranger mouse and time spent in the chamber with the novel object was recorded for 5 min. For the social novelty preference test, mice were returned to the middle chamber, this time with the original mouse (familiar mouse) in its chamber and a new unfamiliar mouse (novel mouse) in the previously empty wire cage. Again, time spent in each outer chamber was recorded for 5 min.

Marble burying test. Mice were placed individually in standard mouse cages containing bedding 5 cm in depth with 20 small marbles arranged in 4 evenly spaced rows of 5 on top of the bedding material. The latency to bury the first marble as well as the number buried (at least 2/3 covered by bedding) was determined in a 30 min trial.

Novel object recognition test. Mice were individually habituated to a 51cm x 51cm x 39cm open field for 5 min. They were then tested with two identical objects placed in the field for 5 min. After two such trials (each separated by 1 minute in a holding cage), the mouse was tested in the object novelty recognition test in which a novel object replaces one of the familiar objects. Behavior was video recorded and then scored for object contact time.

Morris water maze test. Mice were placed into a circular tub filled with opaque water and trained over repeated trials to locate a hidden platform onto which they can sit and escape from the swimming. Each animal received 2 trials per day for 4 days, with a fixed platform location, but a random start position. After being released into the water, each animal was allowed to swim until the platform was found or 90 s had elapsed, at which point the experimenter gently guided the mouse to the platform. A probe trial was given after the completion of training (day 5), in which the platform was removed from the water maze and the animal was allowed to swim freely for 60 s. The amount of time spent in each quadrant was recorded using Noldus Ethovision software and the time spent in the target quadrant (where the escape platform had been) was compared to the average of the times spent in the other 3 quadrants.

Acoustic startle/pre-pulse inhibition (PPI) test. Startle and pre-pulse inhibition testing were performed using San Diego Instruments startle chambers (SR-Lab; San Diego, California). These consist of nonrestrictive Plexiglas cylinders 5 cm in diameter resting on a Plexiglas platform in a ventilated chamber. High frequency speakers mounted 33 cm above the cylinders produce all acoustic stimuli, which are controlled by SR-LAB software. Piezoelectric accelerometers mounted under the cylinders transduce movements of the animal, which are digitized and stored by an interface and computer assembly. Beginning at startling stimulus onset, 65 consecutive 1 ms readings were recorded to obtain the peak amplitude of the animal's startle response. A 60 min test session was used, in which pulse values were 90, 95, 100, 105, 110, 115 and 120 dB; pre-pulse intervals were 25, 50, 100, 200, and 500 ms; and pre-pulse intensities were 78 dB, 82 dB and 86 dB, on a 70-dB background level. Startle pulses were 40 ms in duration and pre-pulses were 20 ms in duration. All trial types (pulse alone, no-stimulus trials (background only), and prepulse + pulse trials) were presented several times in a pseudorandom order (2).

Conditioned fear test. This test enables an assessment of both hippocampus-dependent (contextual portion) and amygdala-dependent (cued portion) learning processes in the same mouse (3,4). During conditioning, mice learn to associate a novel environment (context) and a previously neutral stimulus (conditioned stimulus, a tone and light) with an aversive foot shock stimulus (5). Testing is done in the absence of the aversive stimulus. Conditioned animals, when subsequently exposed to the context or conditioned stimuli with no shock present display the conditioned response, freezing behavior. Freezing behavior in the context and cued tests (relative to the same context prior to shock and an altered context prior to tone, respectively) is indicative of the formation of an association between the particular stimulus (either the environment or the tone) and the shock, indicating that learning has occurred. Conditioning was carried out in Plexiglas Freeze Monitor chambers (Med Associates, Inc.) (26 × 26 × 17 cm) with speakers and lights mounted on two opposite walls, and shockable grid floors. The chambers were housed in sound proofed boxes. Sessions were recorded with real-time digital video and recordings were calibrated to distinguish between subtle movements, such as whisker twitches, tail flicks, and freezing behavior. On day 1, mice were habituated to the chambers in a 5 min shock-free test. On day 2, the mice were subjected to the context and conditioned stimulus (30 seconds, 3000 Hz, 80 dB sound + white light) in association with foot shock (0.60 mA, 2 second, scrambled current). During this 3 min test, the mice received 1 shock, during the last 2 sec of a 30 sec tone/light exposure. On day 3, contextual conditioning (as determined by freezing behavior) was measured during a 5-minute test in the chamber where the mice were trained (context test). The next day, cued conditioning (CS+ test) was tested. The mice were placed in a novel context for 3 minutes, after which they were exposed to the conditioned stimuli (light + tone) for 3 minutes. To create a novel context, the chamber was covered with new walls (white opaque plastic creating a circular compartment in contrast to a clear plastic square compartment) and a new floor (white opaque plastic in contrast to metal grid).
